# Methylene Blue Is a Nonspecific Protein-Protein Interaction Inhibitor with Potential for Repurposing as an Antiviral for COVID-19

**DOI:** 10.1101/2022.03.22.485299

**Authors:** Sung-Ting Chuang, Henrietta Papp, Anett Kuczmog, Rebecca Eells, Jose M. Condor Capcha, Lina A. Shehadeh, Ferenc Jakab, Peter Buchwald

**Author notes:** Corresponding Author, Diabetes Research Institute, Miller School of Medicine, University of Miami, 1450 NW 10 Ave, Miami, FL 33136, USA.

## Abstract

We have previously identified methylene blue, a tricyclic phenothiazine dye approved for clinical use for the treatment of methemoglobinemia and used for other medical applications, as a small-molecule inhibitor of the protein-protein interaction (PPI) between the spike protein of the SARS-CoV-2 coronavirus and ACE2, the first critical step of the attachment and entry of this coronavirus responsible for the COVID-19 pandemic. Here, we show that methylene blue concentration-dependently inhibits this PPI for the spike protein of the original strain as well as for those of variants of concerns such as the D614G mutant and delta (B.1.617.2) with IC_50_ in the low micromolar range (1-5 μM). Methylene blue also showed promiscuous activity and inhibited several other PPIs of viral proteins (e.g., HCoV-NL63 – ACE2, hepatitis C virus E – CD81) as well as others (e.g., IL-2 – IL-2Rα) with similar potency. This non-specificity notwithstanding, methylene blue inhibited the entry of pseudoviruses bearing the spike protein of SARS-CoV-2 in hACE2-expressing host cells both for the original strain and the delta variant. It also blocked SARS-CoV-2 (B.1.5) virus replication in Vero E6 cells with an IC_50_ in the low micromolar range (1.7 μM) when assayed using quantitative PCR of the viral RNA. Thus, while it seems to be a promiscuous PPI inhibitor with low micromolar activity and it has a relatively narrow therapeutic index, methylene blue inhibits entry and replication of SARS-CoV-2, including several of its mutant variants, and has potential as a possible inexpensive, broad-spectrum, orally bioactive small-molecule antiviral for the prevention and treatment of COVID-19.

## INTRODUCTION

Severe acute respiratory syndrome-coronavirus 2 (SARS-CoV-2), a betacoronavirus emerged in late 2019 and responsible for the ongoing COVID-19 pandemic [1–4], is the most infectious agent in a century [5]. Among the seven coronaviruses (CoVs) currently known to infect humans, which include four causing only common colds (HCoV 229E, OC43, NL63, and HKU1) and two that caused previous recent epidemics of high mortality (SARS-CoV-1 ~10% and MERS-CoV ~35%), SARS-CoV-2 caused by far the biggest health and economic damage despite its lower overall mortality compared to the last two. The success of the recent vaccination program notwithstanding, there still is a considerable therapeutic need for antivirals, as a significant portion of the population is unable or unwilling to be vaccinated and mutations are diminishing vaccine efficacy so that SARS-CoV-2 infections are likely to continue in the foreseeable future. Orally bioavailable antivirals are especially of interest since oral treatments can be taken easily following the first symptoms. While efforts toward repurposing of approved drugs SARS-CoV-2 had only very limited success so far [6, 7], two new drugs with classic antiviral mechanisms (i.e., inhibition of protease activity or viral reproduction) have shown promise and granted emergency use authorization by the United States Food and Drug Administration (FDA) for the treatment of COVID-19: molnupiravir (Emory University, Ridgeback Biotherapeutics, and Merck) [8] and nirmatrelvir (Pfizer; as part of the nirmatrelvir/ritonavir combination Paxlovid) [9].

Because of our interest in small-molecule inhibitors (SMIs) of protein-protein interactions (PPIs) [10, 11], we focused on an alternative target and searched for possible inhibitors of the PPIs between the CoV spike (S) proteins and their cognate cell surface receptors needed to initiate cell attachment and virus entry (angiotensin converting enzyme 2, ACE2, for both SARS-CoV-1 and SARS-CoV-2) [12]. While SMIs for PPIs are more challenging to identify than antibodies, progress has been made, and three SMIs of PPIs are now approved for clinical use (venetoclax [13], lifitegrast [14], and fostemsavir [15]) [11, 16–18]. In particular, fostemsavir targets gp120 binding to CD4 to block HIV attachment and entry further validating a PPI inhibitory strategy for antiviral drug discovery. Based on this premise, we initiated a screening to identify possible SMIs of this PPI [12], during which we discovered that methylene blue (MeBlu), a tricyclic small-molecule dye, which is approved for the treatment of acquired methemoglobinemia and some other clinical uses [19–22], inhibits the SARS-CoV-2 – hACE2 PPI and the entry of SARS-CoV-2-spike-bearing pseudoviruses into ACE2 expressing cells [23]. Here, we show that MeBlu inhibits these even for mutants such a D614G and variants of concern (VoC) such as delta (B.1.617.2; first detected in India, October 2020) that have emerged since then. The D614G mutation increases the infectivity, transmission rate, and efficiency of cell entry for SARS-CoV-2 [24] and is present in all later variants of high infectivity. Delta is a VoC that shows increased transmissibility and reduction in neutralization by post-vaccination sera [25]. We also show that MeBlu inhibits the replication of a native/infective SARS-CoV-2 strain (B.1.5) in Vero E6 cells. While we found MeBlu to have apparent promiscuous PPI inhibitory activity and a relatively narrow therapeutic index, this inexpensive and clinically long used drug still could be of interest as an oral treatment for the prevention and therapy of COVID-19, especially in non-industrialized nations with limited access to proprietary vaccines and antivirals.

## MATERIALS AND METHODS

### Binding Assays

Methylene blue and other test compounds used here were obtained from Sigma-Aldrich (St. Louis, MO, USA) and used as such. Purities (and catalog numbers) were as follows: methylene blue >95% (M4159), chloroquine >98.5% (C6628), hydroxychloroquine >98% (H0915), naphthol blue back >99% (70490), and sunset yellow FCF 90% (465224). Proteins used in the binding assays were obtained from Sino Biological (Wayne, PA, USA) as follows: human ACE2-Fc (10108-H05H), SARS-CoV-2 S1 (His tag, 40591-V08H), RBD (His tag, 40592-V08H), D614G S1 (His tag, 40591-V08H3), delta S1 (His tag, 40591-V08H23), delta RBD (His tag, 40592-V08H90), and HCoV-NL63 S1 (His tag, 40600-V08H). HepCV E (His tag, 9146-HC) and CD81-Fc (9144-CD) were from R&D Systems (Minneapolis, MN, USA); IL-2 (His tag, IL2-H52H8) and IL-2Rα-Fc (ILA-H5251) were from ACROBiosystems (Newark, DE, USA). Mutations in the delta S1 variant (40591-V08H23) are T19R, G142D, E156G, 157-158 deletion, L452R, T478K, D614G, and P681R. Mutations in the delta RBD variant (40592-V08H90) are L452R and T478K. Binding inhibition assays were performed in a 96-well cell-free format as described before [12, 23]. Briefly, microtiter plates (Nunc F Maxisorp, 96-well; Thermo Fisher Scientific, Waltham, MA, USA) were coated overnight at 4°C with 100 μL/well of Fc-conjugated receptor proteins diluted in PBS pH 7.2. This was followed by blocking with 200 μL/well of 2% BSA (A7030, Sigma-Aldrich, St. Louis, MO, USA) for 1 h at room temperature. Plates were then washed twice using washing solution (PBS pH 7.4, 0.05% Tween-20) and tapped dry before the addition of the tagged ligand protein and test compounds diluted in binding buffer (20 mM HEPES, pH 7.2) to give a total volume of 100 μL/well. After 1 h incubation, three washes were conducted, and a further 1 h incubation with anti-His HRP conjugate (652504; BioLegend, San Diego, CA, USA) diluted (1:20000) in 2% BSA was used to detect the bound His-tagged ligand. Plates were washed four times before the addition of 100 μL/well of HRP substrate TMB (3,3’,5,5’-tetramethylbenzidine) and kept in the dark for up to 15 min. The reaction was stopped using 20 μL of 1M H2SO4; absorbance values were read at 450 nm. The plated concentrations of Fc-conjugated receptor proteins were 1.0 μg/mL ACE2 for SARS-CoV-2 RBD and D614G S1 variant, 0.5 μg/mL ACE2 for delta RBD and delta S1, 2.0 μg/mL ACE2 for HCoV-NL63, 1.0 μg/mL IL-2Rα for IL-2 and 2.0 μg/mL CD81 for HepCV E. The concentrations of the ligands used in the inhibitory assays were 0.5 μg/mL for RBD, 1.0 μg/mL for D614G S1 variant and delta S1, 0.5 μg/mL for delta RBD, 20 μg/mL for HCoV-NL63, 0.03 μg/mL for IL-2 and 2.0 μg/mL for HepCV E. These values were selected following preliminary testing to optimize response (i.e., to produce a high-enough signal at conditions close to half-maximal response, EC_50_). For all compounds, 10 mM stock solutions in DMSO were used.

### SARS-CoV-2 Pseudovirus Assays

For the BacMam based assays, fluorescent biosensors from Montana Molecular (C1100R and C1110G, C1123G; Bozeman, MT, USA) were used per the instructions of the manufacturer with minor modifications as described before [12, 23]. Mutations in the delta variant (C1123G) are T19R, V70F, T95I, G142D, E156-, F157-, R158G, A222V, W258L, K417N, L452R, T478K, D614G, P681R, and D950N. Briefly, HEK293T cells (CRL-3216; ATCC, Manassas, VA, USA) were seeded onto 96-well plates at a density of 5×10^4^ cells per well in 100 μL complete medium (DMEM supplemented with 10% fetal bovine serum). A transduction mixture containing ACE2 BacMam Red-Reporter virus (1.8×10^8^ VG/mL) and 2 mM sodium butyrate prepared in complete medium was added (50 μL per well) and incubated for 24 h at 37°C and 5% CO_2_. Medium was removed, washed once with PBS, and replaced with 100 μL fresh medium containing test compound at selected concentrations, pre-incubating for 30 min at 37°C and 5% CO_2_. A transduction mixture containing Pseudo SARS-CoV-2 Green-Reporter pseudovirus or Pseudo SARS-CoV-2 Spike Delta Variant Green-Receptor pseudovirus (3.3×10^8^ VG/mL) and 2 mM sodium butyrate prepared in complete medium was added (50 μL per well) and incubated for 48 h at 37°C and 5% CO_2_. Medium was removed, washed once with PBS, replaced with 150 μL fresh medium, and cells incubated for additional 48 h at 37°C and 5% CO_2_. Cell fluorescence was detected using an EVOS FL microscope (Life Technologies, Carlsbad, CA, USA) and was quantified in ImageJ (US National Institutes of Health, Bethesda, MD, USA) [26] using the Analyze Particles tool after thresholding for the corresponding colors.

For the VSV-ΔG based assay, the SARS-CoV-2 S bearing pseudovirus generated in-house was used as described before [12, 27]. Vero-E6 cells (African Green Monkey renal epithelial cells; ATCC cat. no. CRL-1586) engineered to overexpress hACE2/Furin were seeded in 24-well plates to obtain a confluence of 80%. The medium was replaced with 250 μL cell culture medium (DMEM) supplemented with 2% fetal bovine serum, 1% penicillin/streptomycin/glutamine, and the compounds of interest for 30 min at 37°C.

Cells were inoculated with the SARS-CoV-2 spike protein pseudotyped VSV-ΔG (multiplicity of infection = 0.05) by adding complete media to bring the final volume to 400 μL, and 20 h post infection, plates were scanned with a 10× objective using the Incucyte ZOOM imaging system (Sartorius, Ann Arbor, MI, USA). Normalized GFP expression (GCU) values per image were obtained by dividing the Total Green Object Integrated Intensity [Green Calibrated Units (GCU) × μm^2^/image] values of each image by its corresponding Total Phase Area (μm^2^/image) as described before [12, 27].

### Cytotoxicity Assay

For the MTT assay, HEK293T cells were seeded onto 96 wells with the density of 5.0×10^4^ cells per well. Cells were treated with MeBlu for 48h and then replaced with fresh medium for additional 48 h. After that, cells were incubated with MTT reagent at 37°C for 2 h. Remove the medium and dimethyl sulfoxide was added. Cell viability was evaluated using a microplate reader at 570 nm absorbance (SpectraMax iD3, Molecular DEVICES, San Jose, California, USA).

### Surface Plasmon Resonance

SPR was performed at Reaction Biology (Malvern, PA) using single- and multi-cycle kinetics to assess the binding to the SARS-CoV-2 Spike RBD (His-tagged, Arg319-Ph541; RayBiotech) and determine the kinetics/affinity. Measurements were performed on a Biacore 8K (Cytiva) with a series S CM5 (Cytiva) sensor chip. Spike RBD was immobilized via amine coupling at pH 5.5 to immobilization levels of ~1500-2300 RU (single-cycle) and ~2500 RU (multi-cycle). A running buffer of PBS-p+ (20 mM phosphate buffer with 2.7 mM KCl, 137 mM NaCl, and 0.05% surfactant Tween20) with 1% DMSO was used. All data was solvent corrected, reference subtracted, and blank subtracted using the Biacore Insight Evaluation Software. To try and obtain kinetic/affinity parameters, the data was evaluated using a 1:1 kinetic binding model. However, the binding behavior appears to be more complex than can be approximated using a 1:1 binding model, which prevents a quantitative analysis of the SPR binding data.

### *In Vitro* SARS-CoV-2 Antiviral Assay

Assay was performed under BSL-4 conditions [28, 29]. Briefly, Vero E6 cells (European Collection of Authenticated Cell Cultures) were seeded onto 96-well plates at a density of 3×10^4^ cells per well in 100 μL cell culture media that consisted of DMEM (Lonza, Basel, Switzerland), 1% penicillin–streptomycin (Lonza, Basel, Switzerland) and 2% heat-inactivated fetal bovine serum (Gibco, Waltham, MA, USA) the day before the experiment. On the following day, cells were treated with the test compound (MeBlu) at predefined concentrations (14, 12, 8, 6, 4, 2, 1, and 0.5 μM) obtained by diluting DMSO stock solutions with the cell culture media. Immediately after treatment, cells were infected with SARS-CoV-2 (hCoV-19/Hungary/SRC_isolate_2/2020, GISAID ID: EPI_ISL_483637) at MOI: 0.01. Cells were incubated for 30 min at 37°C, then the supernatant was replaced with fresh maintenance media supplemented with the compounds at the appropriate concentration. Nucleic acid extraction was made from the supernatant using a magnetic bead based nucleic acid isolation system (Zybio, Chongqing, China; EXM 3000 Nucleic Acid Isolation System) 48 h post infection. Viral copy numbers were determined using SARS-CoV-2 RdRp gene specific primers and probe (see [28]) and droplet digital PCR (Bio-Rad Laboratories Inc., Hercules, CA, USA; QX200 Droplet Digital PCR System). Copy numbers were normalized to the mean for the untreated, infected wells, *n* = 3 biological replicates. IC_50_ values were determined using non-linear regression analysis (four-parameter logistic model; GraphPad Prism 8).

### Statistics and Data Fitting

Binding inhibition and cell assays were performed in duplicate per plates, and assays were performed as at least two or three independent experiments. As before [12, 23], binding data were converted to percent inhibition and fitted with standard log inhibitor vs. normalized response models [30] using nonlinear regression in GraphPad Prism (GraphPad, La Jolla, CA, USA) to establish half-maximal inhibitory concentrations (IC_50_).

## RESULTS

### Inhibition of SARS-CoV-2 Spike – hACE2 PPIs

We have identified MeBlu as an inhibitor of the SARS-CoV-2 S protein – ACE2 PPI, a PPI essential for the viral attachment and entry of this novel, highly infectious coronavirus [23]. In our ELISA-type assay, MeBlu inhibited the interaction between the receptor binding domain (RBD) of SARS-CoV-2 spike of the original strain and hACE2 in typical concentration responsive manner with an IC_50_ of 3.03 μM (95% CI: 2.54 – 3.61 μM), whereas two other dyes included as control, sunset yellow FCF (FD&C yellow #6; SY) and naphthol blue black (NBlBk), as well as chloroquine showed no such activity (IC_50_ > 500 μM; Figure 1A). In general agreement with this, surface plasmon resonance (SPR) measurements using single- and multicycle kinetics indicated that MeBlu binds to the SARS-CoV-2 spike RBD in the low micromolar range (Supplementary Information, Figure S1). However, the binding behavior appears to be more complex than can be described by a standard 1:1 binding interactions, and since it was unclear from the SPR data alone what model would best describe this behavior, kinetics/affinity values have not been determined. Single-cycle kinetics indicated that MeBlu also binds hACE2 in the low micromolar range, again, in a manner that could not be described well by a standard 1:1 binding interactions (data not shown).

**Figure 1.**
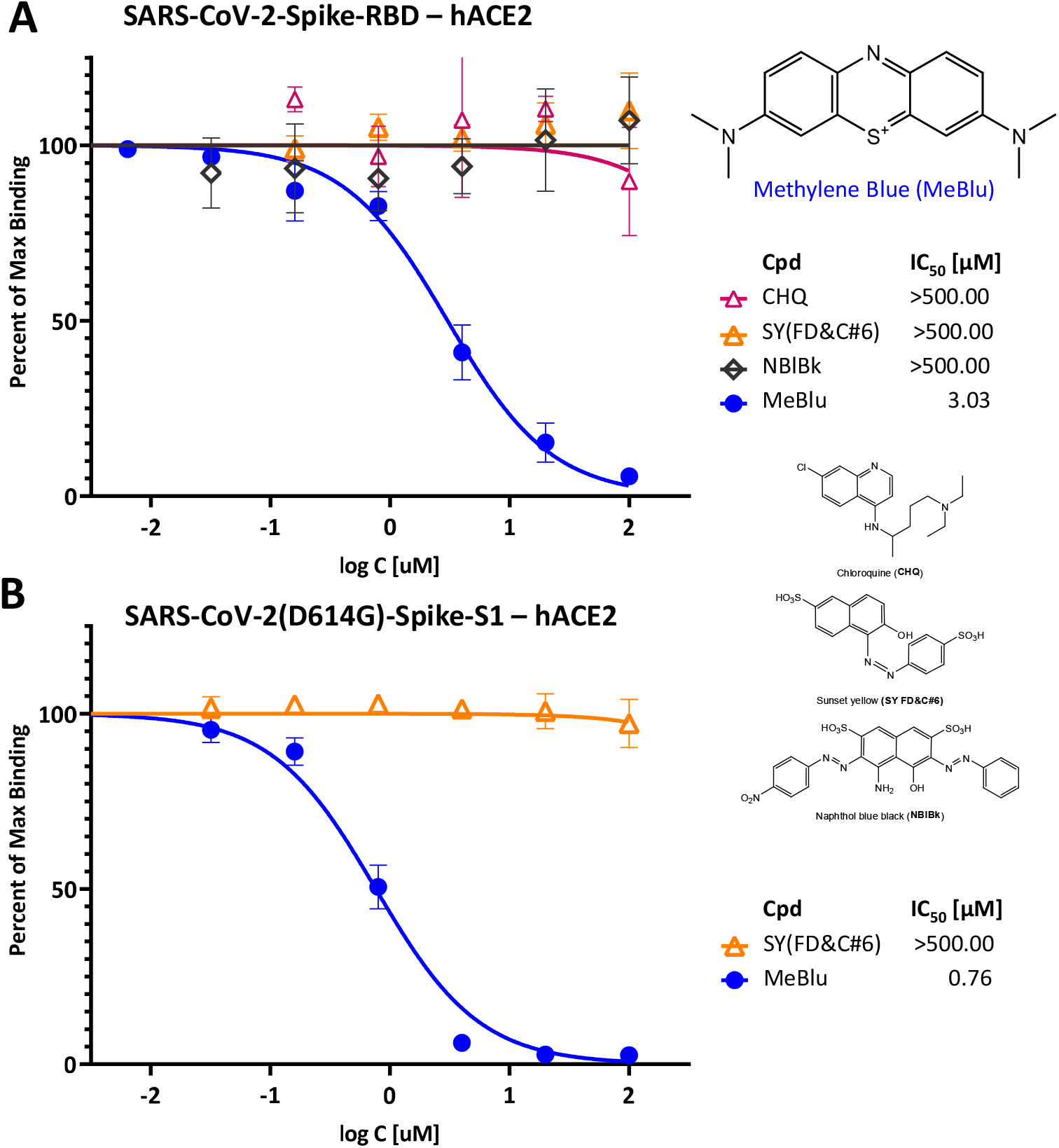
Concentration-dependent inhibition of SARS-CoV-2 Spike protein binding to human ACE2 by methylene blue (MeBlu). Concentration-response curves obtained in ELISA-type assay showing the inhibition of the binding of SARS-CoV-2 spike RBD (His-tagged) (**A**) and D614G mutant S1 (**B**) to hACE2 in the presence of increasing concentrations of test compounds. Contrary to MeBlu, neither chloroquine (CHQ) nor two other dyes included as control (sunset yellow FCF, SY, and naphthol blue black, NBlBk) showed inhibitory activity (chemical structures shown at right). Data (mean ± SD) were normalized and fitted with standard inhibition curves; obtained IC_50_ values are shown.

Notably, especially in light of the emergence of several SARS-CoV-2 mutants and variants of concerns (VoCs) [31], MeBlu also inhibited the binding of mutant spike proteins to hACE2 with very similar activities. For example, MeBlu inhibited binding of the D614G mutant spike protein (S1 portion) with an IC_50_ of 0.76 μM (95% CI: 0.65 – 0.90; Figure 1B). Inhibitory activity remained about the same for the delta (B.1.617.2) variant of concern assayed here either as the S1 protein or as its RBD only: IC_50_s of 1.88 μM (95% CI: 1.47 – 2.39) and 2.42 μM (95% CI: 1.77 – 3.30), respectively (Figure 2) indicating potential for broad-spectrum activity.

**Figure 2.**
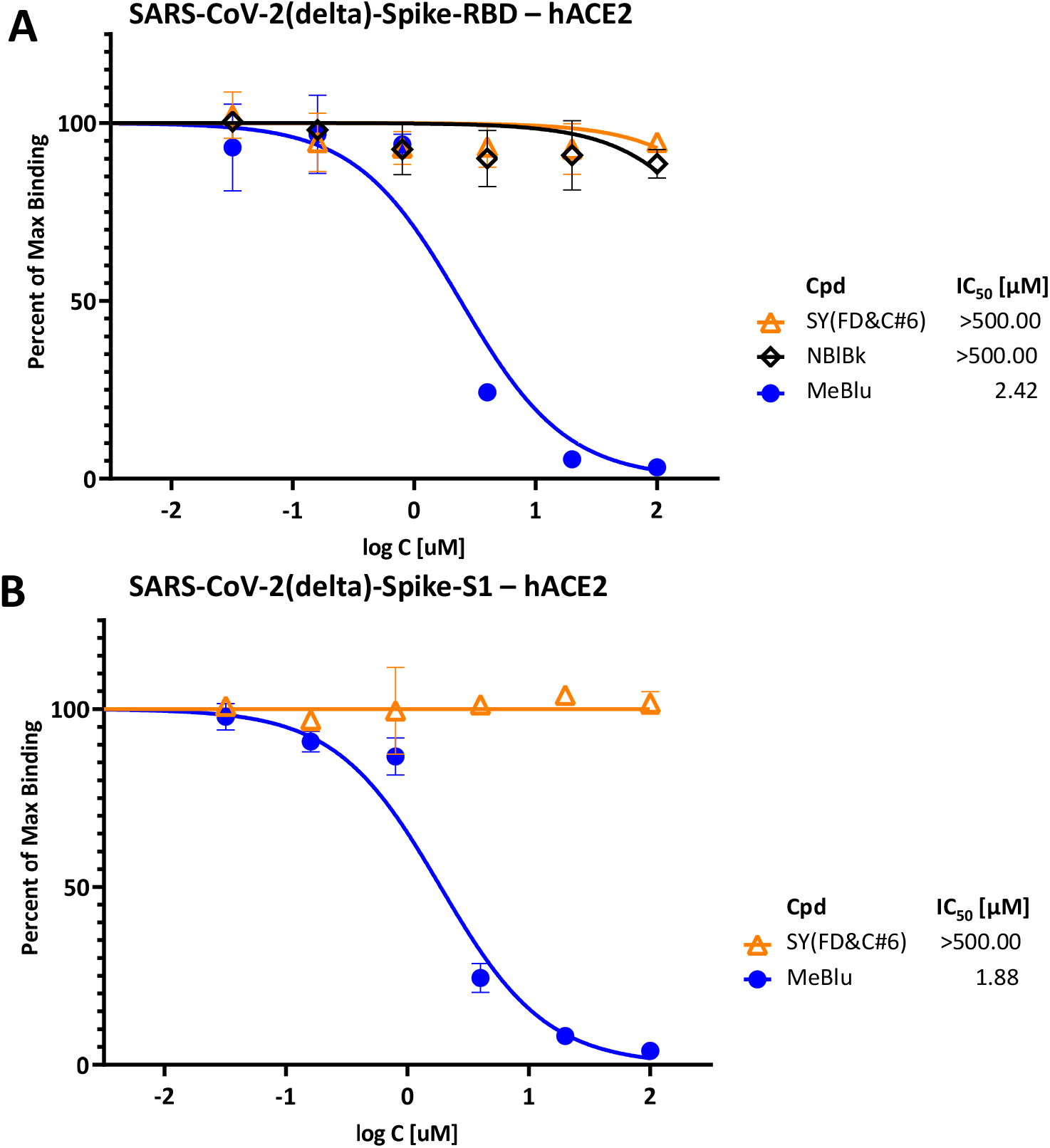
Concentration-dependent inhibition of SARS-CoV-2 delta (B.1.617.2) Spike RBD (A) and S1 protein (B) binding to hACE2 by MeBlu. Concentration-response curves obtained in a similar setup as in Figure 1 but with the spike protein RBD (**A**) and S1 (**B**) of the delta (B.1.617.2) variant in the presence of increasing concentrations of test compounds. Data (mean ± SD) were normalized and fitted with standard inhibition curves; obtained IC_50_ values are shown at right for each figure.

### Inhibition of Other PPIs

It has to be noted, however, that based on our assays, MeBlu seems to be a promiscuous PPI inhibitor with low micromolar activity. It inhibited several PPIs we explored; in fact, it had slightly better activity (lower IC_50_) in a number of them than it had for the SARS-CoV-2 spike – hACE2 PPI. For example, compared to the 3.03 and 2.42 μM for inhibiting the RBD and delta RBD – hACE2 PPIs, respectively, we obtained IC_50_s of 1.93 μM (95% CI: 1.35 – 2.78) for HCoV-NL63 S1 – hACE2, a PPI important for the attachment and entry of this other human coronavirus (HCoV-NL63) that uses ACE2 as its receptor, and 0.80 μM (95% CI: 0.61 – 1.07) for HCV E – CD81, a PPI used by the hepatitis C virus (Figure 3). In addition, MeBlu also inhibited TNF superfamily (TNFSF) PPIs such as TNF-R1-TNFα and CD40–CD154 as mentioned before [23], and the IL-2 – IL-2Rα PPI assayed here (IC_50_ 0.37 μM; 95% CI: 0.27 – 0.50), which we selected for having a complex interface [32] and thus less likely to be susceptible to inhibition by a small molecule.

**Figure 3.**
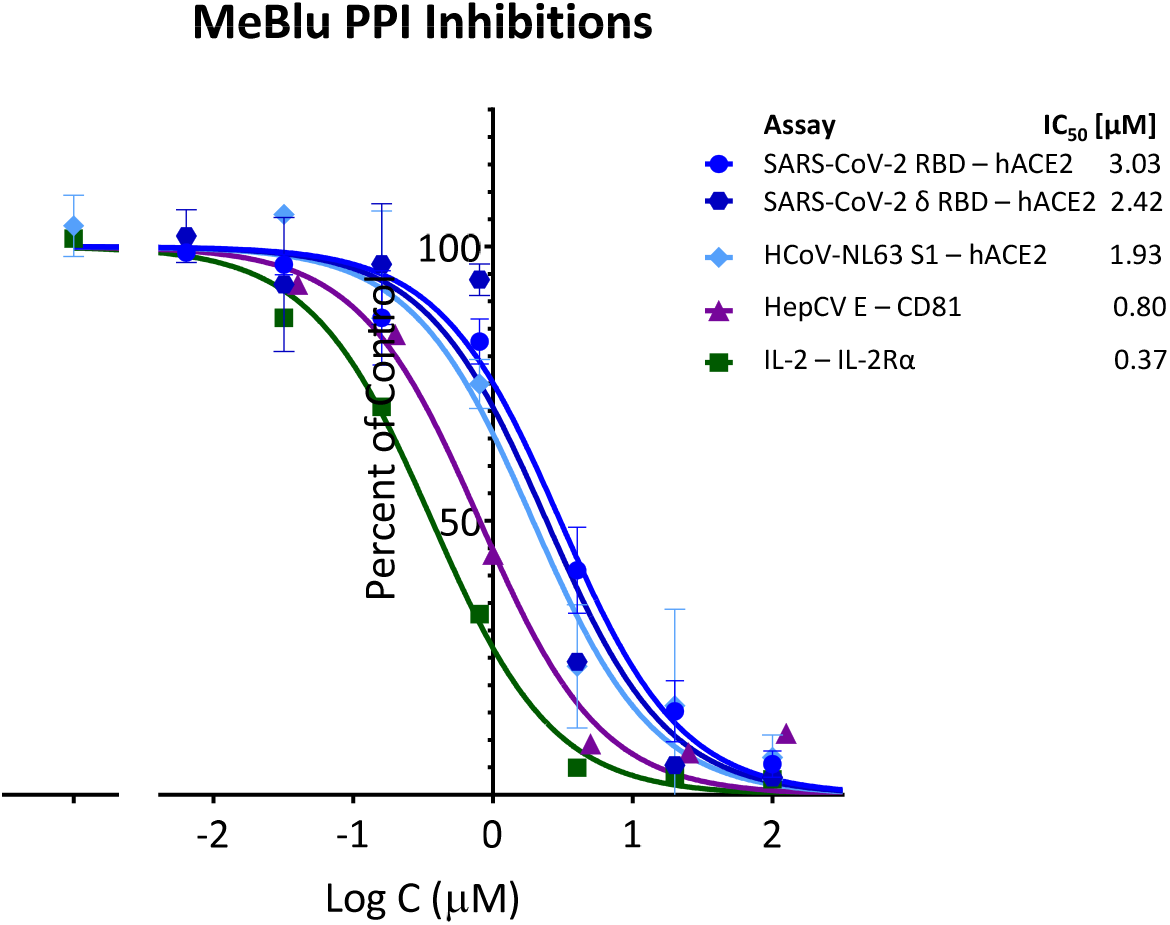
Promiscuous PPI inhibitory activity of MeBlu. Methylene blue inhibited several protein-protein interactions as assayed here in concentration-dependent manner following typical law of mass action response (i.e., unity Hill slope, *n*_H_ = 1). In addition to coronavirus spike protein interactions with human ACE2, including SARS-CoV-2, its delta variant of concern, and the cold causing HCoV-NL63, it also inhibited the PPIs of hepatitis C virus envelope glycoprotein E with CD81 as well as others such as IL-2 – IL-2Rα with similar potency. Data (mean ± SD) for each assay as indicated were normalized and fitted with standard inhibition curves; obtained IC_50_ values are shown at right.

### SARS-CoV-2 Pseudovirus Entry Inhibition

Importantly, this non-specific PPI inhibitory activity notwithstanding, MeBlu also inhibited the entry of two different pseudoviruses bearing the SARS-CoV-2 S spike protein into ACE2-expressing cells. Such pseudovirus assay allows the quantification of viral entry using biosafety level 1 containment because they do not replicate in human cells. First, it has been done with a baculovirus bearing SARS-CoV-2 spike proteins S and generated using BacMam-based tools. As this pseudovirus also expresses bright green fluorescent protein, entry is indicated by expression of green fluorescence in the nucleus of host cells (that express ACE2- and a red fluorescence reporter). If the entry is blocked, the cell nucleus remains dark. MeBlu showed clear concentration-dependent inhibitory activity with IC_50_s of 3.6 μM (95% CI: 2.4 – 5.4 μM) for the original strain and 4.7 μM (95% CI: 3.4 – 6.3) for the delta (B.1.617.2) variant (Figure 4A & B). Here, a set of representative images from serial dilution experiments are also included for illustration (Figure 4C & D). Cytotoxicity evaluation performed under similar conditions gave TC_50_s of 57.9 μM (95% CI: 44.9 – 72.2) indicating clear separation between inhibitory activity and toxicity (Figure 5); nevertheless, MeBlu had a relatively narrow therapeutic (selectivity) index (TI) of just slightly higher than 10 (i.e., a quantitative indicator of the relative safety highlighting the separation between toxic and effective concentrations or doses, e.g., TI = TC_50_/IC_50_, which here in this cell-based assay came out to 15.9 and 12.3 for the original and delta strains, respectively).

**Figure 4.**
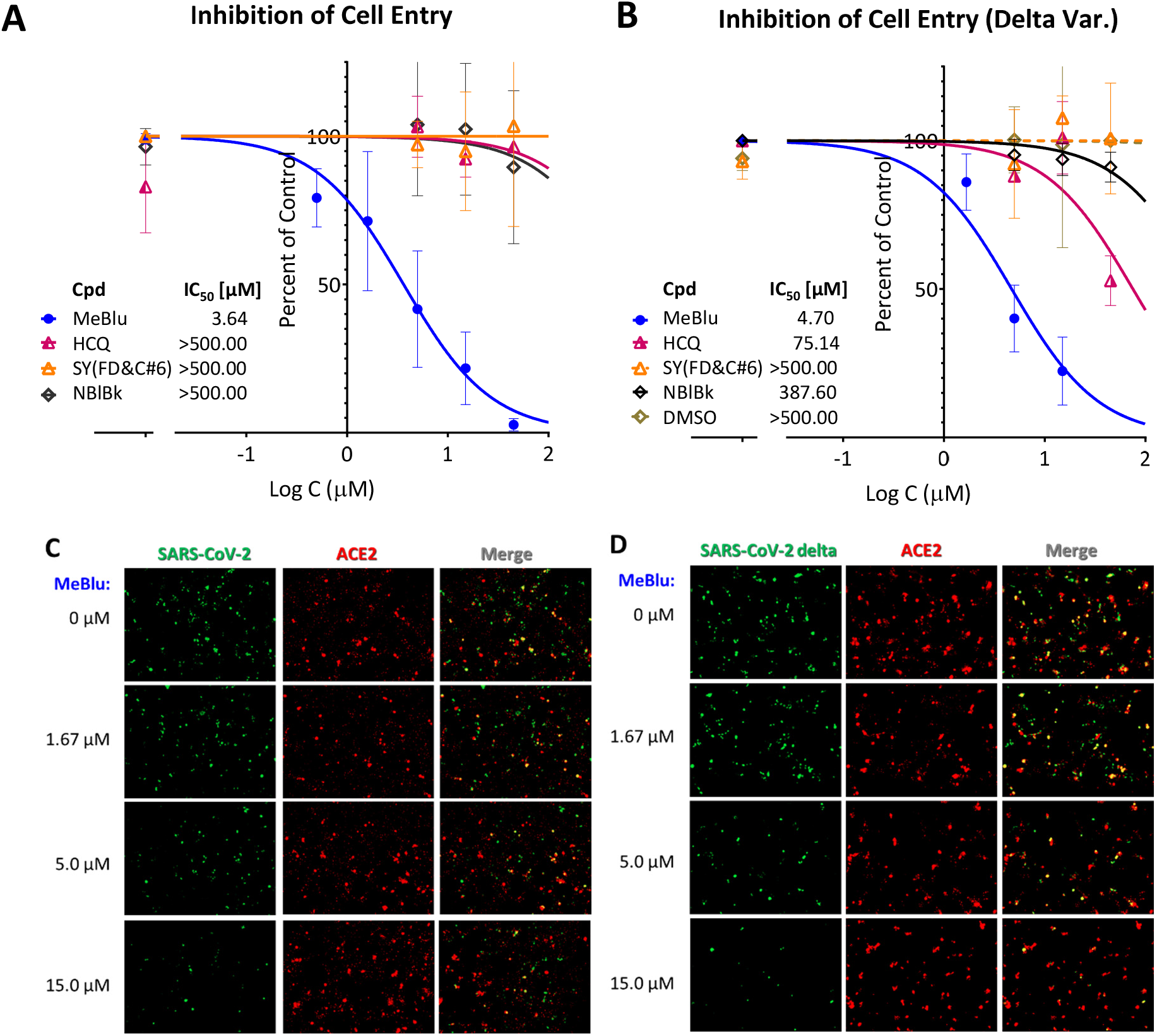
Concentration-dependent inhibition of the entry of SARS-CoV-2 pseudoviruses (BacMam) into ACE2-expressing cells by MeBlu. Quantification of entry of pseudoviruses bearing the original (**A**) or the delta (B.1.617.2) variant (**B**) of the SARS-CoV-2 S protein (plus green fluorescent protein reporters; BacMam-based) in ACE2 (plus red fluorescence) expressing host cells (HEK293T). Values were obtained as before from the quantification of the amount of green present [12, 23], as green fluorescence is expressed only in pseudovirus infected cells. Data are shown as percent of control on semilogarithmic scale and fitted with classic sigmoidal curve used to calculate IC_50_s for MeBlu as well as the control compounds included (hydroxychloroquine, HCQ, sunset yellow, SY, and naphthol blue black, NBlBk). A set of corresponding representative images from serial dilution experiments with MeBlu and the BacMam-based pseudovirus bearing the SARS-CoV-2 S proteins are shown in **C** and **D** for illustration.

**Figure 5.**
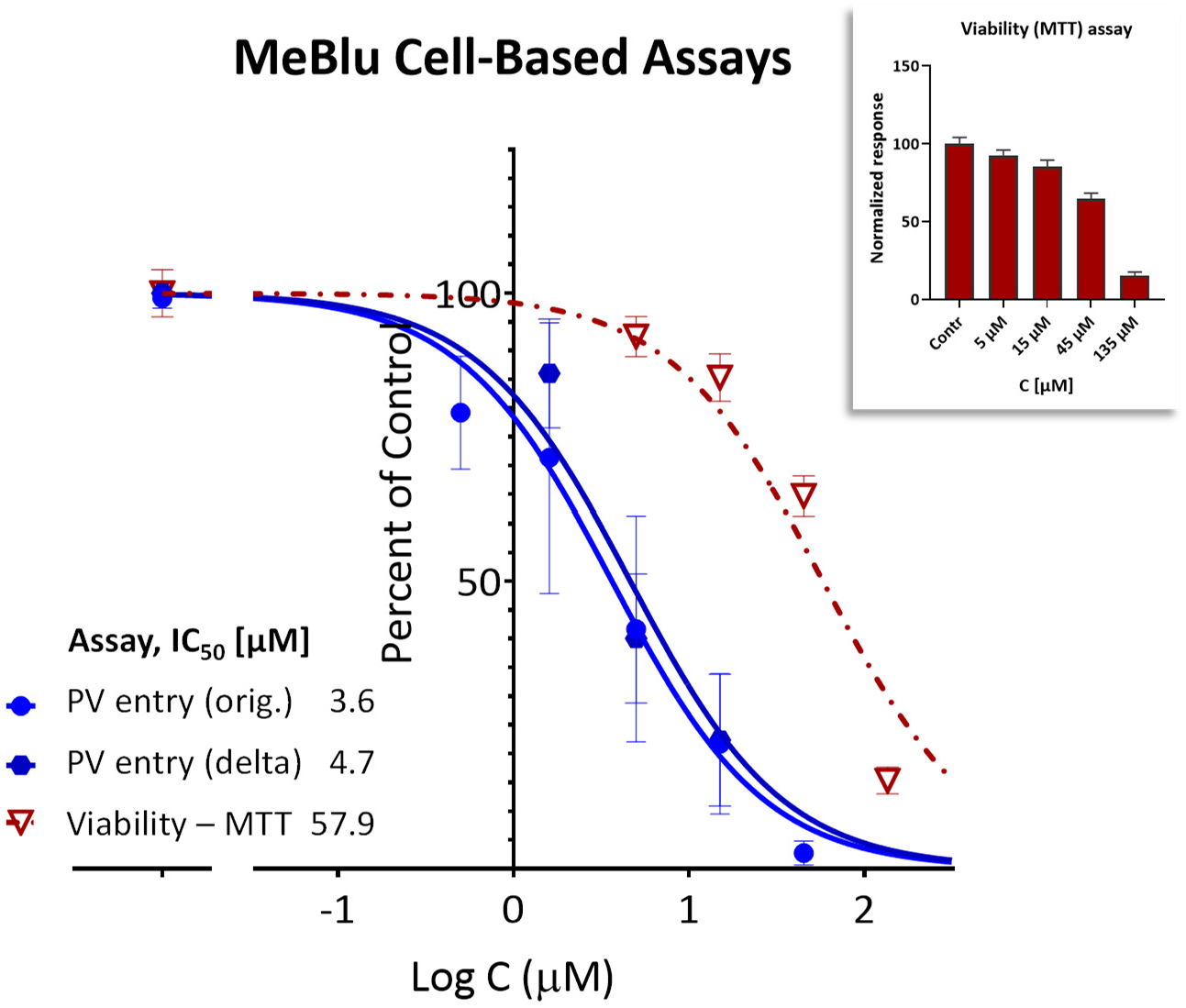
Therapeutic index of MeBlu in the present cell-based assay. Concentration-dependent activity (inhibition of the entry of SARS-CoV-2 BacMam-based pseudoviruses) and safety (effect on viability) of MeBlu in HEK293T cells. Inhibitory effects on the cell entry of pseudoviruses expressing the original and the delta mutant of the spike protein of SARS-CoV-2 (dark and light blue, respectively) shown in parallel with effects on cell viability (dark red; assessed via MTT assay) under the same conditions (48 h in the presence of MeBlu + 48 h culture; also shown separately as a column graph in the inset).

Entry inhibition by MeBlu for the original strain was also confirmed with a different pseudovirus, a SARS-CoV-2 spike plus GFP reporter bearing VSV-ΔG pseudovirus, i.e., vesicular stomatitis virus that lacks the VSV envelope glycoprotein) [27]. As described before [12], GFP fluorescence quantified using a live imaging system (Incucyte) served as a measure of infection and normalized values were fitted with concentration-response curves to obtain IC_50_ (Figure 6). The value obtained in this assay with a different cell line (ACE2/Furin-overexpressing Vero-E6 cells) IC_50_ = 2.5 μM (95% CI: 1.9 – 3.2 μM) was very consistent with that from the previous one confirming the antiviral potential of MeBlu.

**Figure 6.**
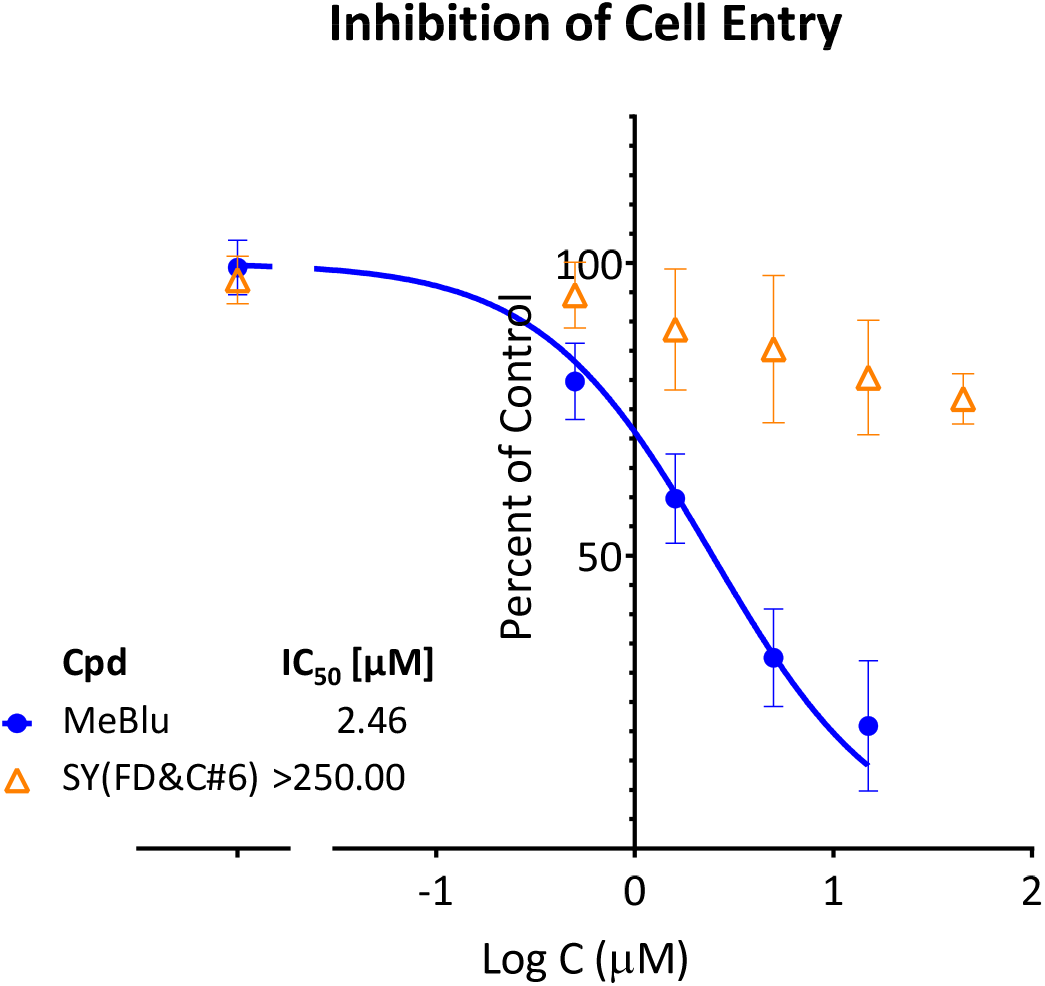
Concentration-dependent inhibition of the entry of SARS-CoV-2 pseudovirus (VSV-ΔG) into hACE2/Furin expressing cells by MeBlu. Entry of VSV-ΔG pseudoviruses bearing the SARS-CoV-2 S protein (plus GFP reporters) in ACE2/Furin overexpressing host cells (Vero E6) in the presence of increasing concentrations of MeBlu was quantified via GFP fluorescence in a live imaging system. Normalized data are shown on semilogarithmic scale and fitted with classic sigmoidal curve used to calculate IC_50_.

### SARS-CoV-2 Antiviral Activity

Finally, the antiviral activity of MeBlu was also confirmed in a viral RNA reduction assay in Vero E6 cells infected with SARS-CoV-2 (B.1.5) at a multiplicity of infection (MOI) of 0.01 as described before [28, 29]. In this assay with live virus, MeBlu was effective in blocking SARS-CoV-2 replication in Vero E6 cells with an IC_50_ of 1.70 μM (95% CI: 1.44 – 1.98) (Figure 7).

**Figure 7.**
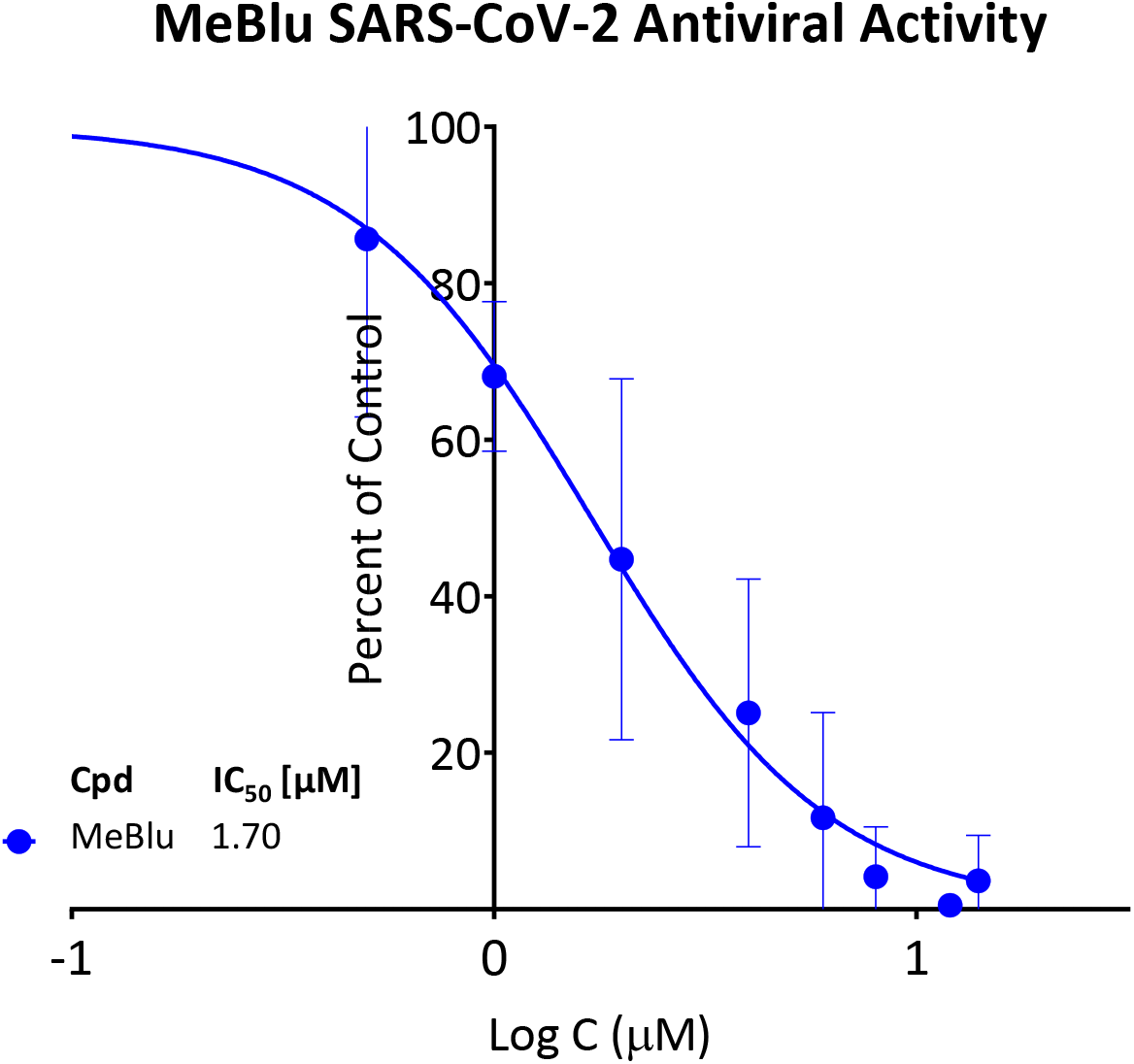
Concentration-dependent inhibition of SARS-CoV-2 replication in Vero E6 cells by MeBlu. Vero E6 cells were infected with SARS-CoV-2 at an MOI of 0.01; treatments were done for 48 h as described in the Methods. The viral yield in cell supernatant was quantified by droplet-digital PCR.

## DISCUSSION

Our results summarized above indicate that MeBlu inhibits the interaction of the SARS-CoV-2 spike protein with hACE2, including for the delta variant of concern (B.1.617.2), and reinforce the potential of MeBlu, a dye compound included in the WHO List of Essential Medicines [19–22] as an inexpensive preventive or therapeutic antiviral treatment for COVID-19. MeBlu, which was first synthesized at BASF (Badische Anilin und Soda Fabrik) by Heinrich Caro in 1876 [21], was, in fact, the first fully synthetic medicine, as it was used for the treatment of malaria since 1891 (until it was replaced by chloroquine during WW2) [20]. MeBlu is approved by the FDA for clinical use in the US for the treatment of methemoglobinemia, and it is also used for some other applications (e.g., prevention of urinary tract infections in elderly patients, treatment of ifosfamide-induced neurotoxicity in cancer patients, treatment of vasoplegic syndrome during coronary procedures, as well as intraoperative visualization of nerves, nerve tissues, and endocrine glands) [20, 22]. In the presence of light, MeBlu has broad-spectrum virucidal activity, and it has been used like this since 1991 to inactivate viruses in blood products prior to transfusions [33]. MeBlu-treated convalescent plasma has also been explored as treatment for COVID-19 patients, e.g., [34]. In addition to our results, others have also found evidence of possible antiviral and, in particular, anti-SARS-CoV-2 activity for MeBlu, even in the absence of UV-induced activation, as summarized in Table 1.

**Table 1.** Summary of published peer-reviewed studies showing anti-SARS-CoV-2 activity for MeBlu.

| Assay | Cell line | IC<sub>50</sub> (μM) | Comment | Reference |
| --- | --- | --- | --- | --- |
| **PPI inhibition (SARS-CoV-2-S – hACE2)** | N/A | 1–5 | Including VoCs (e.g., delta) | [23] and this work (VoC) |
| **SARS-CoV-2 pseudovirus cell entry** | HEK293T | 3–5 | Including VoCs (e.g., delta) | [23] and this work (VoC) |
| **SARS-CoV-2 antiviral activity** | Vero E6 | 1.7 | Inhibition of SARS-CoV-2 (B.1.5) replication | This work |
| **Binding to SARS-CoV-2 S RBD** | N/A | 0.36 | Bio-layer interferometry | [35] |
| **SARS-CoV-2 antiviral activity** | Vero E6 | ~0.3 | Also active against influenza virus H1N1 | [36] |
| **SARS-CoV-2 antiviral activity** | Vero E6 | 0.4 – 1.1 | clinically isolated SARS-CoV-2 strains (IHUMI-3 and IHUMI-6) | [37, 38] |
| **SARS-CoV-2 cytopathicity** | Caco-2 | 2.03 | HTS hit, confirmed in concentration response (Supp Info in Res Square preprint) | [39] |
| **VSV-SARS-CoV-2-Sdel18 pseudovirus entry** | Vero E6 | 9 | HTS hit, confirmed in concentration response | [40] |
| **Image-based multicycle replication assay (hCoV-229E)** | Huh7 | 1.4 | HTS hit, confirmed in concentration response | [41] |

For example, Cagno, Tapparel, and coworkers at the University of Geneva (Geneva, Switzerland) found that MeBlu showed preventive or therapeutic virucidal activity against SARS-CoV-2 (as well as influenza virus H1N1) at low micromolar concentrations and in the absence of UV-activation [36]. Their results also suggested that the antiviral activity of MeBlu might result from multiple mechanisms of action as the extent of genomic RNA degradation was higher in presence of light and after long exposure. Gendrot, Pradines, and coworkers at Institut de Recherche Biomédicale des Armées (Marseille, France) found non-photoactivated MeBlu to inhibit SARS-CoV-2 replication in Vero E6 *in vitro* with an IC_50_ of 0.3 ± 0.03 μM [37] and then confirmed it later with two clinically isolated SARS-CoV-2 strains (IHUMI-3 and IHUMI-6; IC_50_s of 0.4 and 1.1 μM, respectively) [38]. In line with the observations of Cagno and coworkers, this study by Gendrot and coworkers in Vero E6 cells also suggested that MeBlu interacts at both entry and post-entry stages of SARS-CoV-2 infection [38]. MeBlu was also identified in several drug repurposing (repositioning) high throughput screening (HTS) assays looking for anti-SARS-CoV-2 activity, and then confirmed as having low micromolar activity in concentration-response studies. For example, H. L. Xiong and coworkers at Xiamen University (Xiamen, China) found MeBlu to have an IC_50_ of 9 μM in a VSV-SARS-CoV-2-Sdel18 pseudovirus assay when tested as one of the 12 selected compounds of promising activity identified from the screening of 1403 FDA-approved drugs [40]. Finally, an image-based multicycle replication assay based repurposing screening of a chemical library of 5440 compounds that are approved, in clinical trial, or in preclinical development by Murer, Greber and coworkers at the University of Zurich (Zurich, Switzerland) identified MeBlu as one of two promising candidates (the other being mycophenolic acid) that have low micromolar activity against SARS-CoV-2 (1.4 μM). While Cagno et al reported a good therapeutic index for MeBlu in Vero E6 cell-based assay (TI = 46.1/0.11 = 420) [36], this study by Murer et al with Huh7 cells found it to be even lower (TI = 8.7/1.4 = 6.2) [41] than that obtained here with HEK293T cells (TI = 12–16).

Regarding the corresponding clinical applicability of MeBlu, it is important that concentration levels that seem to provide antiviral activity for MeBlu (e.g., IC_50_ = 0.5 – 5.0 μM) are well within the range of those obtained following normal clinical dosage (~3 mg/kg) even with oral administration. For example, 19 μM peak blood concentration were measured after 500 mg p.o. [42], and 6–7 μM trough levels were obtained after doses of 207 mg/day (69 mg, t.i.d., p.o.) [43]. The oral bioavailability *(F* ≈ 80%) and the terminal elimination half-life of MeBlu (*t*_1/2_ ≈ 14 h) [42] are in very reasonable ranges for once daily oral administration. While MeBlu is generally safe, it is known to cause dose-dependent toxicity, in line with the relatively narrow therapeutic index of MeBlu seen here (TI ≈ 10–15; Figure 5). For example, nausea, vomiting, hemolysis, and other undesired side effects start to occur at doses >7 mg/kg (i.e., >500 mg) [19, 21, 22]. MeBlu is contraindicated in persons taking serotonin reuptake inhibitors or having hereditary glucose-6-phosphate dehydrogenase deficiency (G6PD deficiency).

Along these lines, it has to be mentioned that while MeBlu clearly acts as an inhibitor of the SARS-CoV-2 S – hACE2 PPI (Figure 1, Figure 2), it also seems to be a promiscuous PPI inhibitor, which clearly limits its specificity and, hence, usefulness. For example, it also inhibited TNFSF PPIs (e.g., CD40–CD40L, TNF-R1–TNFα) [23] as well as the IL–2R–IL-2 PPIs in our assays with similar low micromolar potency (Figure 3). The three-ring phenothiazine framework of MeBlu resembles the three-ring framework of erythrosine B, an FDA-approved food colorant that we have shown before to act as promiscuous PPI inhibitor (together with some other structurally similar xanthene dyes such as rose Bengal and phloxine) [44]. Furthermore, MeBlu is well-known to show polypharmacology and act on a multitude of targets [20]. While these further limit specificity and can cause unwanted effects, at least some of these effects could actually be beneficial in treating COVID-19 [45, 46]. For example, its main mechanism of action behind its clinical use is its ability to reduce the oxidized form of hemoglobin when in a state of methemoglobinemia and increase the oxygen-binding capacity of hemoglobin, which increases oxygen delivery to tissues [22]. This could be an important additional benefit in COVID-19 patients who often exhibit very low oxygen levels, i.e., silent (or happy) hypoxemia [47].

There is some, albeit limited clinical evidence supporting the use of MeBlu to prevent or treat COVID-19, and most of it should be treated with caution as it was published not in internationally well-recognized peer-reviewed medical journals. The first support for a possible preventive role came from a retrospective study of 2500 patients treated with MeBlu (75 mg t.i.d.) as part of their cancer care in France. This found that none of them developed influenza like illness during the early phases of the COVID-19 pandemic (March 2020) [48]. Currently, there are three trials documented as having been initiated with MeBlu for COVID-19 treatment (according to ClinicalTrial.gov): one in Mexico (NCT04619290; 1 mL Prexablu and 50 min low-level light therapy s.i.d. for 7 days), one in Switzerland (NCT04635605; MeBlu 100 mg b.i.d. for 5 days), and one in Iran (NCT04370288; MeBlu 1 mg/kg as part of a three-drug last therapeutic option add-on cocktail with vitamin C 1500 mg/kg and N-acetyl cysteine 2000 mg/kg in critically ill COVID-19 patients). The last one completed Phase 2 and 3 studies with promising results claimed. In Phase 1, four of the five treated patients responded well to the treatment [49]. The Phase 2 extension was a randomized, controlled, open-label clinical trial involving 80 hospitalized patients with severe COVID-19. It indicated that addition of MeBlu to the treatment protocol significantly improved oxygen saturation (SpO_2_) and respiratory distress (*p* < 0.001) resulting in decreased hospital stay (7.3_±4.7_ vs. 11.7_±6.6_ days, *p* = 0.004) and mortality (12.5% vs. 22.5%) [50]. A Phase 3 randomized, controlled, openlabel clinical trial was completed in 223 hospitalized patients with confirmed severe cases of COVID-19. This also found that addition of oral MeBlu to the standard care significantly shortened hospital stay (6.2_±3.1_ vs 10.6_±9.2_, *p* < 0.001), and decreased mortality (12.2% vs 21.4%, *p* = 0.07) [51].

Intravenous MeBlu (1 mg/kg) has been explored as a rescue therapy in 50 moderate to severe hypoxic COVID-19 patients with acute respiratory distress syndrome (ARDS) with promising results and no major side effect or adverse events [52]. Finally, inhaled applications have also been explored in some developing countries as respiratory treatments [53]. Two studies with nebulized MeBlu conducted in patients with COVID-19 infections in India claimed noticeable benefits: decrease in inflammatory markers and oxygen requirements and a trend toward reduced hospital stays (9.17 vs 12 days) in an observational study with 63 patients divided into three groups [54] and beneficial effects on oxygen saturation and requirements in 7 patients with severe COVID-19 and ARDS who did not respond to antiviral (remedesivir, favipiravir) and anti-inflammatory (tocilizumab, steroids, heparin) treatments and oxygen inhalation, but subsequently responded to MeBlu inhalation [55].

## CONCLUSION

In conclusion, we found that MeBlu is a low-micromolar inhibitor of the PPI between SARS-CoV-2 spike and its cognate receptor ACE2 including for mutants such as the delta variant of concern. MeBlu also inhibited the cell entry of spike-bearing pseudoviruses both for the original strain and the delta variant, and it blocked SARS-CoV-2 (B.1.5.) virus replication in Vero E6 cells. Thus, MeBlu, a drug approved for clinical use, shows potential as a repurposed (repositioned) antiviral agent for the prevention and treatment of COVID-19. While MeBlu shows strong polypharmacology, seems to be a nonspecific PPI inhibitor, and has a relatively narrow therapeutic index, it still has potential as an inexpensive, widely available, and orally administrable treatment for diseases caused by SARS-CoV-2, its variants of concern, and possibly other spike-expressing CoVs.

## ABBREVIATIONS

The following abbreviations are used in this manuscript:

ACE2: angiotensin converting enzyme 2
CoV: coronavirus
MeBlu: methylene blue
NBlBk: naphthol blue black
PPI: protein-protein interaction
SARS: severe acute respiratory syndrome
SMI: small-molecule inhibitor
SPR: surface plasmon resonance
SY: sunset yellow FCF
TNF: tumor necrosis factor
VoC: variant of concern.

## ACKNOWLEDGEMENTS

We are grateful to Dr. György Miklós Keserű for his help facilitating the collaboration with the National Laboratory of Virology group at the University of Pécs, Hungary.

## SUPPLEMENTARY MATERIALS

Supplementary material for this article is available at the end of this file and includes Figure S1, Binding of MeBlu to SARS-CoV-2 spike RBD as assessed via SPR.

## AUTHOR CONTRIBUTIONS

STC performed the binding and BacMam pseudovirus assays and analyzed the corresponding data; HP performed the live virus assays and analyzed the corresponding data; AK performed the ddPCR assay; RE performed the SPR assays, analyzed the corresponding data, and contributed materials; JMCC performed the VSV pseudovirus assays; LAS provided materials and analyzed the data; FJ provided materials and supervised the anti-SARS-CoV-2 experiments; PB conceived and designed the project, provided study guidance, analyzed the data, and wrote the draft manuscript. All authors contributed to writing, have read, and agreed to the published version of the manuscript.

## FUNDING

Financial support by the Diabetes Research Institute Foundation (www.diabetesresearch.org) is gratefully acknowledged (PB). The anti-SARS-CoV-2 antiviral assay was financed by the Thematic Excellence Program 2021 Health and National Defense, National Security Subprograms of the Ministry for Innovation and Technology in Hungary within the framework of the EGA-10 and NVA-07 projects of the University of Pécs and the Comprehensive Development for Implementing Smart Specialization Strategies at the University of Pécs (EFOP-3.6.1.-16-2016-00004) grants. Dr. Shehadeh is funded by grants from the National Institute of Health (1R01HL140468) and the Miami Heart Research Institute.

## INSTITUTIONAL REVIEW BOARD STATEMENT

Not applicable.

## INFORMED CONSENT STATEMENT

Not applicable.

## DATA AVAILABILITY STATEMENT

The raw data supporting the conclusions of this manuscript will be made available by the authors upon reasonable requests from qualified researchers.

## CONFLICTS OF INTEREST

The authors declare no conflict of interest.

## SUPPLEMENTARY INFORMATION

### Supplementary Figures

**Figure S1.**
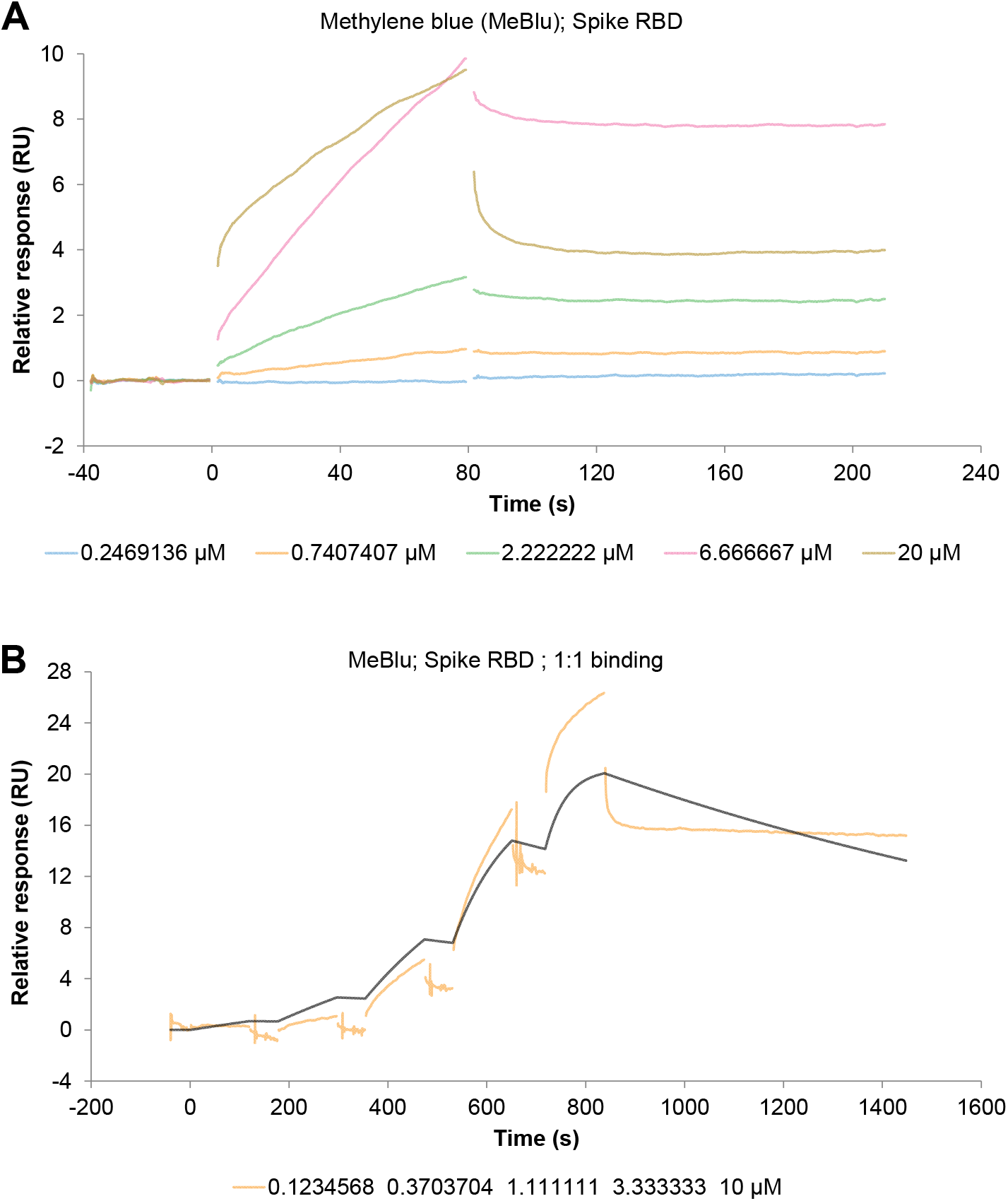
Binding of MeBlu to SARS-CoV-2 spike RBD as assessed via surface plasmon resonance (SPR). Multi-(**A**) and single-(**B**) cycle kinetics were used to measure binding to the SARS-CoV-2 spike RBD using a Biacore 8K instrument with a series S CM5 sensor chip (Cytiva) as described in the Methods. Initially a multi-cycle kinetic measurement was performed, however, the signals clearly did not return to baseline over the dissociation period and a regeneration condition was not established to ensure removal of all previously bound molecule before the injection of the next concentration. As a result, there were a decreasing number of binding sites available over the course of the measurement as sites became blocked by the preceding concentration(s), which meant this approach was not appropriate to use for a quantitative analysis even though qualitatively it was clear that binding was occurring. To deal with the apparent slow dissociation (signal not returning to baseline between injections) a single cycle kinetic approach was used instead. Data indicate low-micromolar binding; however, the binding behavior could not be described well with a 1:1 binding interactions, thus preventing a quantitative analysis to determine the on/off-rate and/or KD values. Figure B shows an example of this deviation from 1:1 binding where the colored (yellow) curve is the measured data while the black curve is the fit curve produced using a 1:1 binding model. Instead, the SPR data provides qualitative support that MeBlu interacts with SARS-CoV-2 spike RBD.

## Notes

### Competing Interest Statement

The authors have declared no competing interest.

